# Molecular address codes delineate circuit-specific motor neuron-muscle matchmaking in the developing tetrapod limb

**DOI:** 10.64898/2026.08.04.742930

**Authors:** Fabio Sacher, Bianka Berki, Maëva Luxey, Antoine Fages, Patrick Tschopp

## Abstract

An essential part of nervous systems development is the establishment and refinement of correct circuit architectures. In vertebrate neuromuscular circuits, this requires connecting the axons of motor neurons in the central nervous system to their corresponding muscle groups in the periphery. This process can roughly be sub-divided into two distinct phases: axon guidance towards their innervation territories, followed by the establishment of correct nerve-muscle connections. While the role of attractive and repulsive cues in axon guidance has been studied extensively, relatively little is known about the molecular interactions shaping motor neuron to muscle matchmaking that contribute to circuit specificity and refinement.

Here, using the tetrapod limb as a model system, we focus on the maturation of three distinct neuromuscular circuits, targeting different proximal-distal and anterior-posterior territories of the developing chick forelimb, and sequence the transcriptomes of individually backfilled motor neurons as well as of the muscles they connect to. Comparative analyses across circuits suggest the presence of distinct molecular “address codes”, based on individual signatures of matching profiles of secreted signaling factors and cell surface receptors. Furthermore, we probe for inherent transcriptional plasticity in this system, using an experimentally altered limb periphery that results in nerve miswiring, and sequence the corresponding motor neuron-muscle pairs. Lastly, to facilitate data exploration, we present an R Shiny app, to investigate circuit-specific ligand-receptor interaction profiles in both neuron-muscle and muscle-neuron directions, in control and experimentally altered limb configurations.

Collectively, we present a resource to investigate the cellular and molecular basis for muscle-nerve matching during neuromuscular circuit refinement in the tetrapod limb, with implications for our understanding of vertebrate neuromuscular development and evolution, as well as regenerative approaches targeting re-innervation after injury or nerve damage.

## Introduction

In most animal species, active locomotion is essential for procuring food, finding mates, evading predation, and range expansion by migration. In tetrapods, the land-dwelling vertebrates and their descendants, this predominantly relies on the coordinated movement of their paired fore- and hindlimbs, with morphological variations therein often reflecting adaptations to distinct modes of locomotion (Coates 1994; Shubin, Tabin, and Carroll 1997; Rothier et al. 2024). While the general limb architecture is largely dictated by the underlying skeleton, full functionality depends on its proper integration with nerves and muscles, in form of distinct neuromuscular circuits. Correct circuit assembly is thus a crucial step during limb development. As the skeletal progenitors condense, muscle precursors delaminate from the somitic dermomyotome, migrate into the limb bud and form dorsal and ventral muscle masses, before splitting into individual muscles and eventually connecting to the correct bones *via* tendons (Kardon 1998; Schweitzer, Zelzer, and Volk 2010). Concomitantly, motor neurons project their axons from the lateral motor columns (LMCs) in the spinal cord, at fore- and hindlimb levels, to innervate the peripheral limb muscles, with molecularly distinct pools of motor neurons foreshadowing muscle-target specificity (Bonanomi and Pfaff 2010; Stifani 2014; Dasen 2022). In addition, proper integration of sensory neurons and interneurons completes circuit architecture and functionality (Plant, Weinrich, and Kaltschmidt 2018).

For motor neurons, proper circuit development relies on axon guidance mechanisms, as growth cones explore the limb periphery to find the appropriate muscle territories in the limb periphery. Depending on the repertoire of expressed receptors on an axonal growth cone, a neuron can react to repulsive or attractive guidance cues, which can either be acting at long ranges for secreted factors, or locally, for membrane bound factors (Huber et al. 2003; Russell and Bashaw 2018). Limb-innervating motor axons project through the anterior part of the sclerotome, and coalesce to form the brachial and lumbosacral plexi, at the base of the fore- and hindlimbs, respectively (Wang and Anderson 1997; Roffers-Agarwal and Gammill 2009). There, outgrowth pauses and axons reshuffle, before forming distinct nerve bundles that project into either the dorsal or ventral part of the limb (Bonanomi 2019). For example, dorsal attraction is mediated by GDNF (glial cell line-derived neurotrophic factor), present at this decision point and perceived by susceptible neurons expressing *Ret*, while ventrally expressed repulsive cues help to ensure appropriate projection patterns (Kramer et al. 2006; Bonanomi et al. 2012). From the plexii outwards, nerves continue to grow and split in a stereotypical and highly reproducible fashion, to project to the different limb quadrants where their cognate muscle targets form (Hirasawa and Kuratani 2018; Bonanomi 2019; Luxey et al. 2020). However, much less is known about the molecular nature of pathfinding decisions at these more distal limb locations. Upon reaching their target muscles, nerves defasciculate and form neuromuscular junctions at the location of acetylcholine receptor clusters, before being ensheathed by terminal Schwann cells, which contribute to maturation and synapse elimination (Darabid, Perez-Gonzalez, and Robitaille 2014; Li, Xiong, and Mei 2018). Neurons which fail to establish functional connections with their muscles retract and die, due to limiting amounts of muscle-derived neurotrophic factors important for motor neuron maturation and survival (Henderson et al. 1994; Haase et al. 2002). However, molecular cues that potentially fine-tune this motor neuron-muscle matchmaking process, during initial contact and circuit-specific maturation, have yet to be identified.

To address this crucial gap in our understanding, here we capitalize on retrograde axonal backfilling and single-cell RNA-sequencing of individual motor neurons and combine it with muscle-specific bulk RNA transcriptomes, to define circuit-specific molecular address codes in the developing tetrapod limb. In particular, we investigate the transcriptional profiles of motor neurons and their corresponding muscles in the context of three distinct neuromuscular circuits, innervating different wing territories along the proximal-distal and anterior-posterior axes of the chicken wing. Supplementing our data with samples from an experimentally altered periphery, we probe for inherent transcriptional plasticity in the system, in response to supernumerary muscles and resulting nerve miswirings in a model of induced polydactyly. Finally, we package our data for user-friendly exploration of ligand-receptor interactions in form of a publicly available dockerized R Shiny app.

## Results

### Circuit-specific molecular signatures in chick forelimb motor neurons and muscles

To investigate the molecular basis of motor neuron-muscle matchmaking, we investigated three distinct neuromuscular circuits in the developing chicken forelimb, at embryonic day 9, i.e., Hamburger-Hamilton stage 35 (HH35) (Hamburger and Hamilton 1951). We probed transcriptomes at the level of their motor neurons and muscles, in control and polydactyl configurations. Circuits were selected to represent different proximal-distal and anterior-posterior muscle target locations in the chicken forelimb – specifically, circuits involving the proximal-posterior muscle *Flexor carpi ulnaris* (FCU), the distal-posterior muscle *Flexor digiti quarti* (FDQ), and the distal-anterior muscle *Abductor medius* (AM) (Fig. 1A). We focused our attention on the ventral side of the limb, and the FDQ circuit in particular, to capitalize upon the previously reported plasticity in nerve and muscle patterns occurring there in an experimentally induced model of polydactyly: this manipulation eventually results in a splitting of the Median nerve, and its partial miswiring towards a duplicated FDQ muscle (see (Luxey et al. 2020), and red arrow in Fig. 2A).

**Figure 1.**
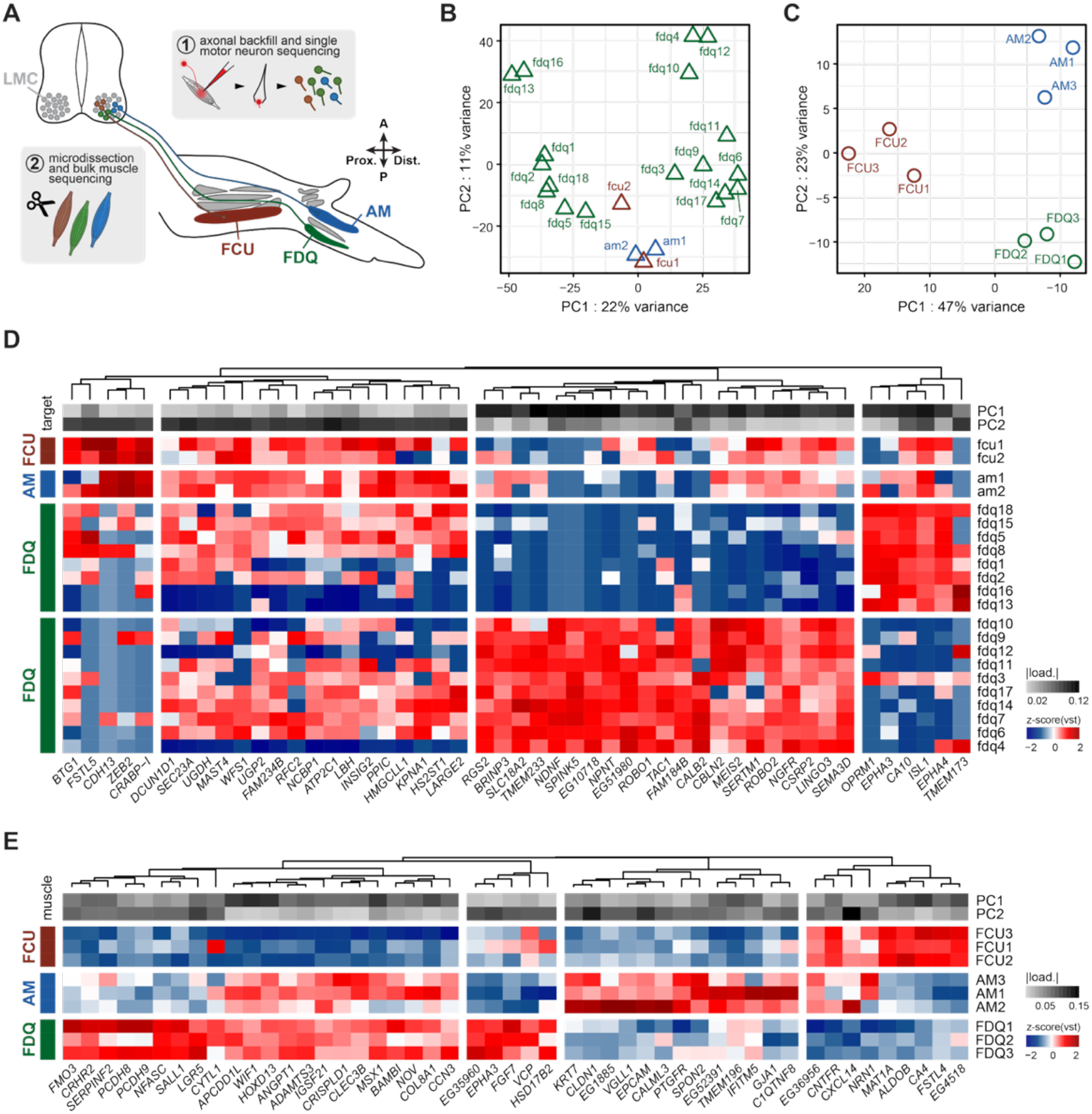
Transcriptomic profiles of distal limb muscles and their innervating motor neurons. **(A)** Sampling scheme for isolating control motor neurons via retrograde axonal backfill, and micro-dissecting the corresponding muscle targets. **(B-C)** Principal component analysis (PCA) of expression profiles of control neurons and control muscles based on the top 500 variable genes. Motor neurons are colored by their target muscle. **(D-E)** Heatmap of the intersection of the 25 genes with highest absolute loading values for PC1 and2. Expression values are based on variance-stabilizing transformation of the top 500 variable genes, and row clustering is based on Pearson correlations. Greyscale shows the absolute loading values for PC1 and 2. Colors correspond to **A**. PC = principal component; PCA = principal component analysis; vst = variance stabilizing transformation. Targeted circuits by muscle are ‘fcu’-FCU, ‘fdq’-FDQ and ‘am’-AM.

**Figure 2.**
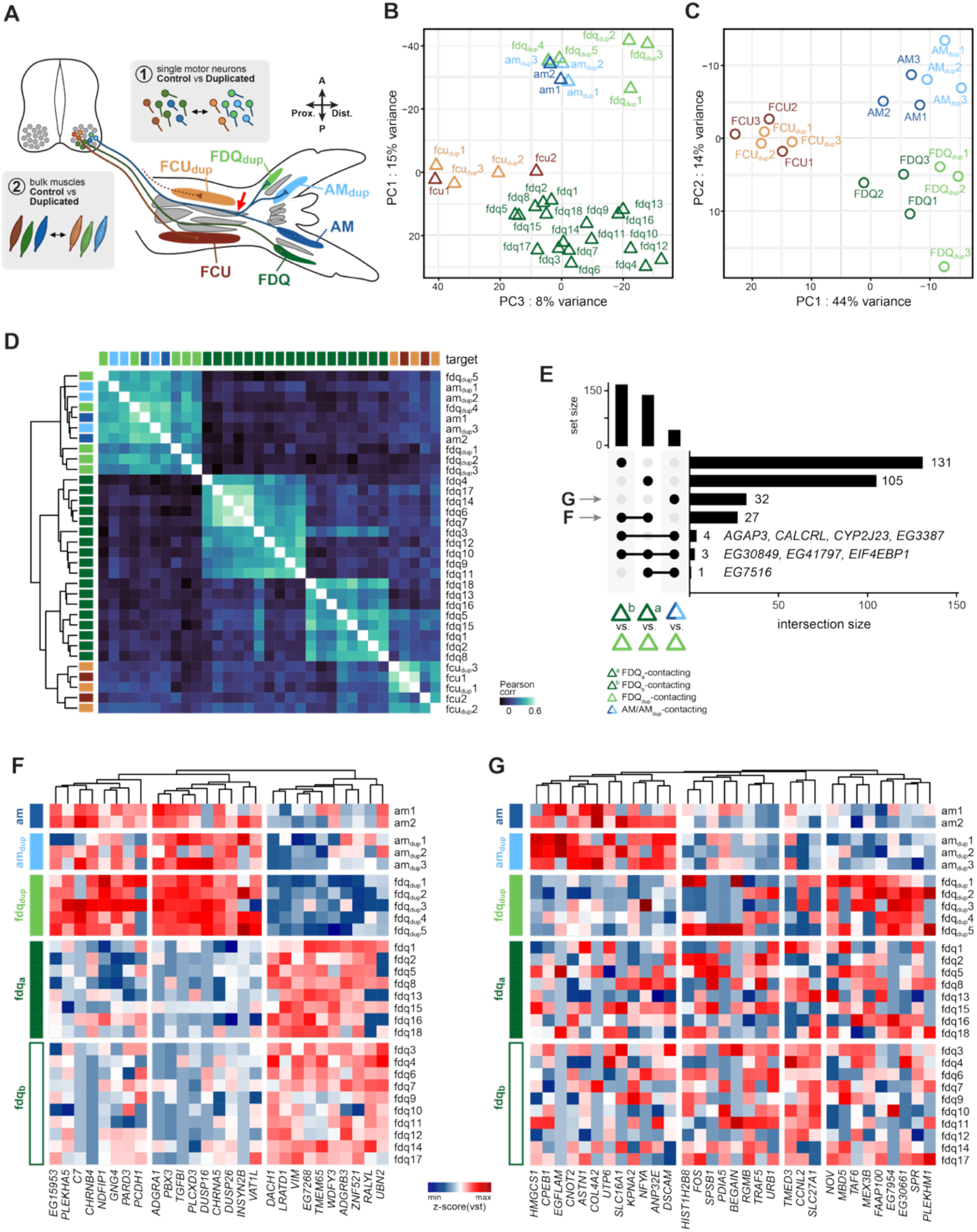
Changes in transcriptomes upon distal muscle duplication in an induced polydactyly. **(A)** Sampling scheme for isolating duplicated motor neurons via retrograde axonal backfill, and micro-dissecting the corresponding muscle targets. Red arrow indicates the branching point of the Median nerve, innervating AM_dup_ and FDQ_dup_ muscles in the duplicated ‘posterior’ side. **(B)** Principal components PC1 and PC3 of a principal component analysis (PCA) of all neurons based on the top 500 most variable genes. Axes are arranged to align samples with the position of the target muscles. Colors correspond to **A. (C)** PCA of all muscles based on the top 500 variable genes. Colors correspond to **A. (D)** Heatmap of pairwise Pearson correlation values across all neurons, based on variance stabilizing transformation (vst) expression of the top 500 variable genes. Samples are colored by their target muscle, corresponding to **A. (E)** Upset plot showing the intersections of differentially expressed (DE) genes between ‘fdq_dup_’ neurons and ‘am_dup_’, ‘fdq_a_’, or ‘fdq_b_’ neurons. Arrowheads indicate the comparisons plotted in **F** and **G. (F-G)** Heatmap of z-scored vst expression values for the DE genes between ‘am_dup_’ and ‘fdq_dup_’ **(F)** and ‘fdq_a_’ and ‘fdq_b_’ with ‘fdq_dup_’ neurons **(G)**. Columns clustered by Euclidean distance. Rows are ordered by target muscle.

For sequencing of individual motor neurons transcriptomes, we employed an in-house developed method for muscle-specific retrograde axonal backfill coupled with spinal cord tissue dissociation and picking of single labeled neurons (see Materials & Methods and (Berki et al. 2023) for details). For corresponding muscle transcriptomes, we micro-dissected entire muscles and extracted total RNA, followed by Smart-Seq2 library preparation and sequencing (see Materials & Methods). In total, we collected 33 single motor neuron cell bodies and 18 muscle samples. Throughout our analyses motor neuron samples are referred to by lowercase labels reflecting the muscle (labeled in uppercase) they were connected to (e.g., ‘fcu’ motor neuron samples were collected from FCU muscle backfills). Reads were mapped to the chicken *Galgal6* genome assembly and further analyzed using an in-house custom pipeline (see Fig. S1A,B; for sequencing depth and gene detection rates, please refer to Supplementary Tables S1 and S2). Motor neuron cell type identity was verified by marker gene expression profiles, including the pan-neuronal marker *TUBB3*, genes involved in acetylcholine signaling (*CHAT, COLQ, SLC18A3*), lateral motor column (LMC) specification (*ALDH1A2* and *FOXP1*) and vesicle trafficking and exocytosis (*CPLX1* and *SNCA*) (Delile et al. 2019; Sacher et al. 2026). Additionally, we checked for absence of markers for sensory neurons – namely, *NTRK1, NGF*, and *RUNX3 –*, to exclude potential cellular contaminants in our sampling from the dorsal root ganglia cells (Faure et al. 2020; de León, Gibon, and Barker 2021), as they can be labelled by retrograde backfills as well (Fig. S1A). Muscle samples expressed low levels of early lineage markers (*MYF5, PAX3*, and *SIX4*), but elevated levels of later myogenic factors (*MYOD1* and *MYOG*), myosins (*MYL1, MYL2*, and *MYL10*), and troponins (TNNC1, TNNC2, and TNNI2) (Fig. S1B) (Sutherland et al. 1993; Grifone et al. 2005).

We first focused our analyses on motor neuron and muscle transcriptomes coming from the three circuits in their standard configurations (Fig.1A). Principal component analysis (PCA) of gene expression in neuronal samples was dominated by our ‘oversampling’ of FDQ-innervating motor neurons: namely, along principal component 1 (PC1, 22% of variance), two distinct sub-groups of FDQ-innervating neurons were identified – hereafter referred to as ‘fdq_a_’ and ‘fdq_b_’ –, with ‘fcu’ and ‘am’ samples clustering together (Fig. 1B). Differential expression (DE) analyses between the two ‘fdq’ sub-groups revealed several potential marker genes, including transcriptional regulators and axon guidance molecules (Fig. S1C,F). While GO-term enrichment analyses for both sets of DE genes failed to reveal any clear functional differentiation (Fig. S1D), in ‘fdq_b_’ neurons, several embryonic marker genes associated with the innervation of fast-twitch muscle fibers appeared enriched, such as *PCP4, CALB2* or *MAFB* (Fig. S1C,F) (D’Elia et al. 2023a). Unexpectedly, the majority of these neurons did not express *ISL1* – a canonical marker of the medial LMC (LMC_m_) that innervates the ventral limb – and five cells instead showed trace amounts of lateral LMC (LMC_l_) markers *LHX1* and *ETV4* (Lee et al. 2023) (Fig. S1E). This hinted at a possible LMC_l_ fate for these cells, potentially due to accidental mistargeting while collecting the samples. However, given the care taken during the injection procedures, and the prevalence of this molecular signature (9 out of 18 ‘fdq’ cells total), we favor the ‘fast-*versus* slow-twitch fiber’ distinction as a possible explanation for the observed transcriptional differences. From the PCA of gene expression of muscle samples, a clearer image emerged: replicate samples clustered in a muscle-specific manner, reflecting their relative spatial arrangements inside the intact limb. There was a clear separation between the proximal FCU and the two distal FDQ and AM muscles along PC1 (47% of variance), as well as an anterior-posterior differentiation of all three groups along PC2 (23% of variance, Fig. 1C).

To inspect the genes driving the observed groupings in the two PCAs, we plotted unsupervised hierarchically clustered expression heatmaps for the top 25 loading genes, for principal components 1 and 2, in control motor neuron and muscle samples (Fig 1D,E). As expected, genes driving PC1 in our motor neuron PCA largely separated ‘fdq_a_’ from ‘fdq_b_’ neurons. Among these genes, we identified several neurotrophic factors, axon guidance molecules and transcription factors (Fig 1D, top). Along PC2, *CRABP-I*, a retinoic acid binding protein, the transcription factor *ZEB2*, and the atypical cadherin *CHD13* were major contributors to the observed variance and showed particularly high expression in ‘fcu’ and ‘am’ neurons (Fig 1D, bottom). For muscle samples, top PC1-loading genes reflected the general architecture of the proximal-distal limb axis. For example, the proximal FCU muscles lacked *HOXD13* expression, a marker of the distal limb, which in turn showed robust expression in the two distal muscles, the AM and FDQ (Fig 1G) (Dollé et al. 1991; Nelson et al. 1996). More intriguingly, however, we were able to identify distinct molecular identities for all three muscles along PC2, partially delineated by members of the FGF and BMP signaling pathways, as well as by the expression profiles of extracellular matrix (ECM) proteins and genes involved in axon guidance (Fig 1G).

Collectively, by profiling transcriptomes of paired motor neuron and muscles samples, we uncover evidence for distinct and circuit-specific molecular signatures in both entities that may play instrumental roles during the establishment and refinement of neuromuscular networks in the tetrapod limb.

### Transcriptional responses upon peripheral muscle duplications and distal nerve miswiring

To probe for transcriptional plasticity in the system, upon the experimental induction of extra peripheral muscle targets, we next focused on our motor neuron and muscle samples coming from polydactyl forelimbs. Polydactylies were induced by implanting retinoic acid (RA)-soaked beads in the anterior wing margin at embryonic day 3 (HH19), which induces complete mirror duplications at zeugopod and autopod levels, both in the skeleton and at the level of the musculature (Fig. 2A, see Materials & Methods) (Tickle et al. 1982; Duprez et al. 1999; Luxey et al. 2020). Importantly, upon such duplications, the global innervation pattern changes for one of our targeted muscle pairs. Namely, the duplicated posterior and distal muscle FDQ_dup_ is now innervated by a split Median nerve, unlike its counterpart FDQ that still connects with the Ulnar branch as in control limbs (Fig. 2A, red arrow). Both AM and AM_dup_, as well as FCU and FCU_dup_ maintain the respective connections to their major nerve branches (Luxey et al. 2020).

This anatomically apparent rewiring was also reflected in the respective motor neuron transcriptomes. Through PCA, we identified three clusters of control and duplicated motor neurons along PC1 and 3, with ‘fcu’/’fcu_dup_’ and ‘fdq’ neurons forming two of the main clusters. Neurons connecting to the AM, AM_dup_, and FDQ_dup_ muscles, however, grouped together, indicating a close transcriptional similarity between neurons belonging to these three experimental groups (Fig. 2B). Along PC2, again a split between control ‘fdq_a_’ and ‘fdq_b_’ neurons was apparent (Fig. S3A). In contrast, muscle control and duplicate pairs always clustered together, despite the change in innervation for the FDQ/FDQ_dup_ pair (Fig. 2C). Hence, not only was the relative positioning and morphology of duplicate muscles preserved, but also their respective molecular identities appeared to be largely maintained (Luxey et al. 2020). Differential expression (DE) analyses between controls and duplicates, revealed the highest number of DE genes in the distally located FDQ/FDQ_dup_ muscle pair, followed by AM/AM_dup_ and only minor differences in FCU/FCU_dup_ (Fig. S2A-D). Expression differences in duplicated muscles seemed to reflect a slight delay in maturation, with several ECM components showing elevated expression levels in controls (Fig. S2B-D) (Sacher et al. 2021; Wilde et al. 2021). Moreover, we found *MYL2, MYL3*, and *MYOZ2* downregulated in duplicated AM and FDQ muscles, indicating a reduction of slow muscle fibers that we confirmed with immunohistochemistry against MF20 (muscle specific myosin heavy chain) and NA8 (anti-slow myosin heavy chain 2, slow twitch muscle fibers) (Fig. S2B,C and S2E-H) (Reiser, Greaser, and Moss 1988).

To further investigate possible transcriptional consequences of the observed neuronal miswiring – i.e., motor neurons belonging to the Median nerve now contacting the FDQ_dup_ muscle – we first calculated Pearson correlation values on the variance stabilizing transformed (vst) expression levels of the top 500 variable genes in all our motor neuron samples, followed by unsupervised hierarchical clustering. Ulnar nerve cells targeting the FCU, FCU_dup_ and FDQ muscles formed one major cluster, with muscle- and subgroup-specific sub-divisions, while motor neurons contacting the AM, AM_dup_ and FDQ_dup_ muscles formed the second major transcriptional clade (Fig. 2D). We then performed differential expression analyses, between ‘fdq_dup_’ and ‘am/am_dup_’ samples – i.e., cells belonging to the same transcriptional clade and nerve branch, but contacting molecularly and morphologically distinct muscles –, as well as ‘fdq’ motor neurons in control configuration (Supplementary Table S3). Most DE genes were identified in comparisons between ‘fdq_dup_’ and ‘fdq’ samples, in line with these cells contributing to two distinct nerve branches, the Median and Ulnar, respectively (Fig. 2E, Fig. S3C-E). Likewise, genes showing consistent expression differences between the two ‘fdq’ control populations and ‘fdq_dup_’, revealed a clear ‘am’-like signature in ‘fdq_dup_’ cells (Fig. 2F and Fig. S3D,E). Intriguingly, however, genes that were differentially expressed between ‘am’ and ‘fdq_dup_’ showed a more ‘fdq’-like behavior in ‘fdq_dup_’ cells, indicating that the mis-wiring of a Median-nerve motor neuron can result in the partial transcriptional reprogramming towards an FDQ-like muscle target identity (Fig. 2G).

### Interrogating paired ligand-receptor expression profiles in a circuit-specific manner

The transcriptional response of rerouted ‘fdq_dup_’ cells – i.e., of motor neurons contributing towards the Median nerve whilst connecting to a duplicated FDQ muscle – to a more ‘fdq’-like state suggested the presence of a muscle-to-neuron feedback, likely mediated by the circuit-specific secretion and perception of distinct signaling molecules. Indeed, previous work has shown that GDNF, an important neuronal survival cue, is specifically expressed in the *Latissimus dorsi* and *Cutaneus maximus* muscles at later stages of development, and plays an instrumental role in motor neuron positioning and maturation (Haase et al. 2002). Accordingly, we next focused our attention on the expression dynamics of known ligands and receptors, and their communication potential based on a curated database of human cell-cell signaling interactions (Ramilowski et al. 2015). We sub-setted these lists for chicken orthologs and first performed PCA of all expressed ligands or receptors in motor neuron and muscle samples. Interestingly, these PCAs, based only on the expression profiles of signaling ligands and receptors, recovered the anatomical arrangements of all our samples (Fig. S4A-D). We therefore concluded that functionally distinct motor neurons and muscles of the developing tetrapod limb are molecularly delineated, potentially in a circuit-specific manner. Amongst the top 30 most variably expressed ligands and receptors from our samples, we identified multiple candidates that showed either motor neuron- or muscle group-specificity. Furthermore, several putative signaling interactions were identified, in both muscle-to-neuron and neuron-to-muscle direction, as suggested by the curated ligand-receptor interaction database and indicated in grey lines connecting the two pairs of heatmaps (Fig. 3A,B and C,D). To investigate such putative signaling interactions between corresponding neuron-muscle pairs in a more systematic and targeted manner, we defined an “interaction score”, as the product of average ligand and receptor Transcript Per Million (TPM) expression values in physically connected motor neurons and muscles (Fig. 4A). Additionally, to test for the degree of circuit-specificity in these interactions, we statistically compared the interaction scores of correct interactions – i.e., between physically connected neurons and muscle pairs – *versus* incorrect interactions – i.e., between motor neuron and muscle samples with no known physical contact in the intact limb (see Material & Methods). We then ordered the interactions by motor neuron-muscle groups and p-values and selected the top interactions for each circuit for visualization in a dot plot-like manner (Fig. 4B). To facilitate data exploration, we packaged our interaction analysis results in an interactive and dockerized R Shiny app. Across 33 motor neuron and 19 muscle samples, in control and polydactyl configurations, the app allows for the targeted inspection of 2,253 direction-sensitive interaction scores for the investigated circuits, i.e., 1,123 from neuron to muscle and 1,130 from muscle-to neuron (Fig. 4C). The list of all interactions is searchable by circuit logic – at the level of the motor neurons or corresponding muscles –, as well as ligands or receptors, in a fully reciprocal manner. In a typical query, the user can first screen interaction scores for e.g. circuit specificity (Fig. 4B, see arrow), interrogate to what extent these scores are driven by either ligand or receptor expression levels, in motor neurons or muscles (Fig. 4D), and then explore potential molecular ambiguities with alternative receptors (Fig. 4D, top) or ligand partners (Fig. 4D, bottom). All resulting plots are easily exported, for downstream analyses and presentation purposes, and the app with its associated datasets is publicly available on *github*.

**Figure 3.**
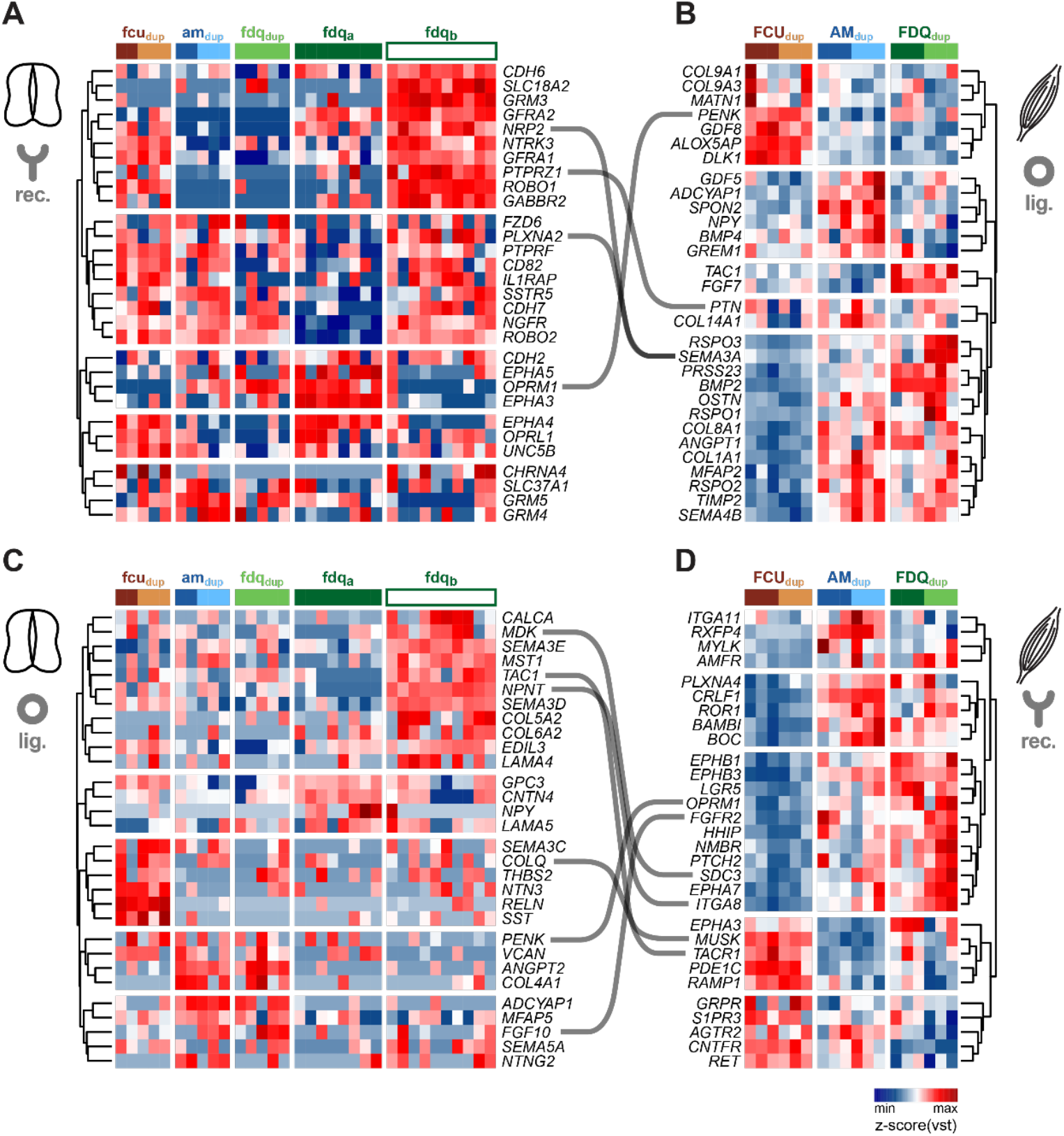
Ligand Receptor interactions between neurons and muscles. **(A-D)** Heatmaps of z-scored vst expression values of the top 30 most variable expressed ligands and receptors across all neuron and muscle samples. Rows clustered on Euclidean distances. Grey lines indicate potential interactions of corresponding ligand and receptor pairs, as reported in Ramilowski et al., 2015. **(A-B)** represents muscle ligand-neuron receptor direction. **(C-D)** represents neuron ligand-muscle receptor direction.

**Figure 4.**
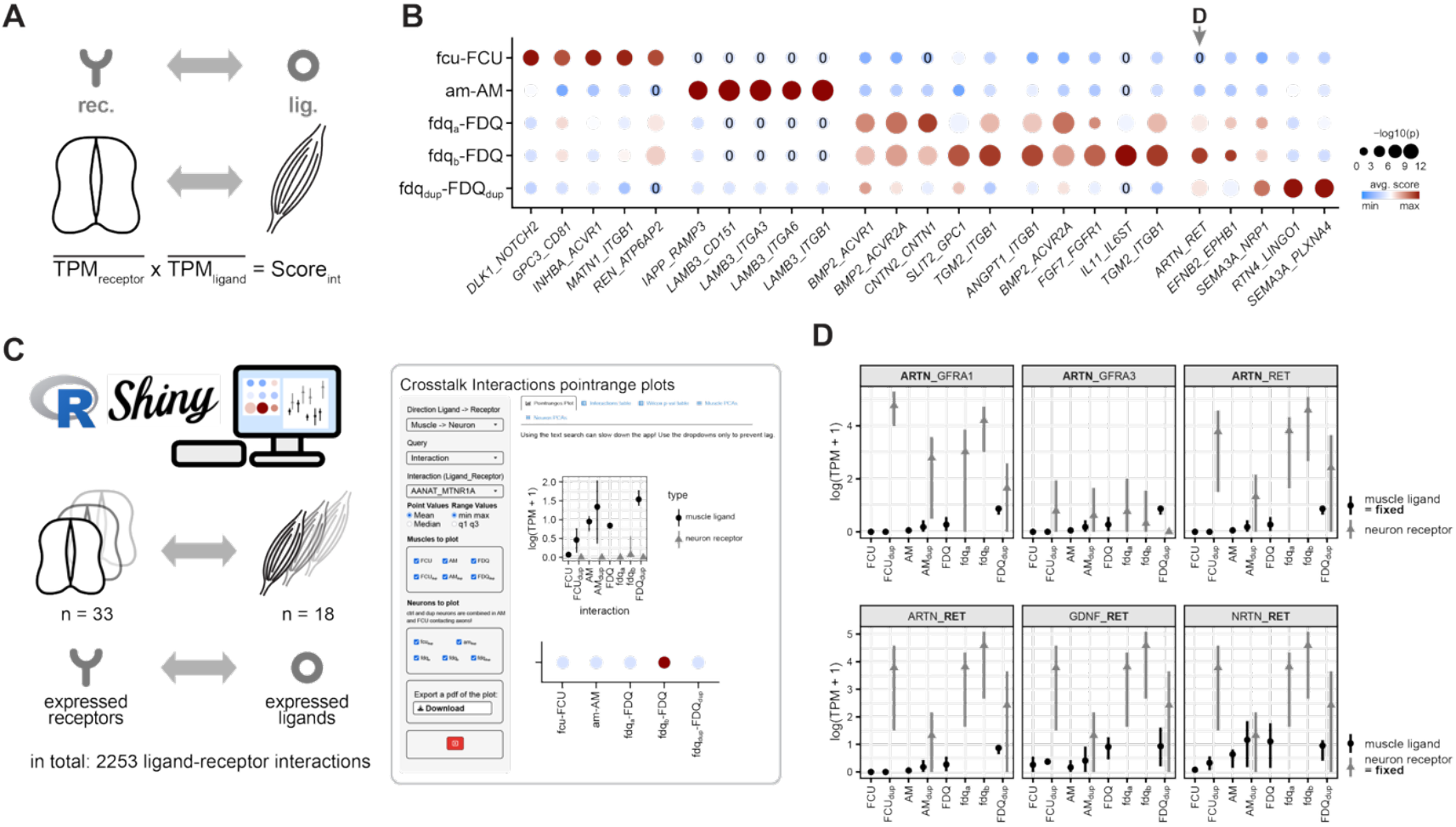
Exploring ligand-receptor interactions with an interactive R Shiny app. **(A)** Scheme describing the logic for calculating ligand-receptor interaction scores, in paired motor neurons and their muscle targets. **(B)** Dotplot showing the top 5 interactions by p-value per motor neuron receptor-muscle ligand interaction. Dot size represents -log_10_(p-value) and the colors represent z-scored average interaction scores. P-values are defined by comparing correct interactions vs incorrect interactions (see Material and Methods for details). Arrow indicates candidate interactions highlighted in **D. (C)** Outline of the R Shiny app. It contains ligand-receptor interaction scores in both directions, i.e. neurons to muscles and muscle to neurons, as well as expression ranges for select ligands and receptors in a circuit-specific manner. **(D)** Example point ranges plots for muscle ligand-motor neuron receptor interactions, here focusing on iterations and alternative interaction partners for *ARTN* (ligand) and *RET* (receptor).

## Discussion

In tetrapod vertebrates, the correct assembly and maturation of limb neuromuscular circuits is critical for coordinating the contraction patterns of their peripheral musculature, and hence controlled locomotion in adult life. While the molecular mechanisms governing the early embryonic pathfinding decisions in these circuits have been extensively studied (Huber et al. 2003; Bonanomi and Pfaff 2010; Russell and Bashaw 2018), considerably less is known about the molecular basis of target matching and circuit refinement once motor axons reach the developing musculature. The recent rise in cellular throughput and transcript detection sensitivity in various single-cell transcriptomic approaches now offers appropriate tools for comprehensive molecular profiling of neuronal diversity (Shekhar et al. 2016; Armand et al. 2021), including for the cholinergic neuron sub-class essential for muscle innervation (Alkaslasi et al. 2021). Here, capitalizing on targeted high-coverage single cell transcriptomes of backfill-labeled and handpicked motor neurons, combined with corresponding bulk muscle profiling, we define circuit-specific molecular address codes active during this maturation process. We identify restricted ligand-receptor interaction pairs, in motor neurons and muscles, suggestive of distinct molecular avenues for cross-tissue communication and circuit maturation during tetrapod limb development. Furthermore, by probing rewired circuits in a model of experimentally induced polydactyly, we demonstrate the plastic nature of the system with potential implications for the evolution of novel morphologies and locomotory functions in the tetrapod limb, as well as regenerative approaches aiming at muscle-nerve rewiring after injury or disease.

### Circuit-specific molecular address codes in motor neuron-muscle pairs

The existence of pool- and, hence, potentially muscle-specific molecular subtypes in LMC motor neurons has been extensively documented, early during tetrapod limb development, as well as in isolated cases at postnatal stages (Dasen et al. 2005; Kramer et al. 2006; Stifani 2014; Mendelsohn, Dasen, and Jessell 2017). Here we probe the intervening maturation stages, during prehatching development, at single-cell resolution. In motor neurons, we identify distinct combinations of transcription factors, secreted ligands and cell surface receptors, ECM components, and axon guidance molecules that were associated with circuit-specific neuronal transcriptomes. For example, transcriptional co-factors – such as *MEIS1, MEIS2* or *PBX3* – show sub-group-specific transcriptional signatures and positive expression correlations with a previously identified motor neuron maturation gene co-expression module, as does *NR0B1*, a transcriptional repressor of the nuclear receptor gene family so far only implicated in hypothalamus development (Fig. 1D, Supplementary Table S5 and Supplementary Table S6) (Zhang et al. 2022; Bobola and Sagerström 2024; Sacher et al. 2026). Circuit-specific transcription factor codes and multi-protein complexes could thus provide a gene regulatory logic underlying the divergent effector gene trajectories we observe during motor neuron maturation (Arendt et al. 2016; Wang et al. 2025; Sacher et al. 2026). These expression profiles also support a model in which neuron-to-muscle matchmaking is eventually refined through combinatorial molecular address codes, rather than a small number of universal recognition factors, down to the fiber-specific subtype (Dasen 2017; D’Elia et al. 2023b). Lastly, selective expression of trophic factors in distinct neuronal subsets, such as *NDNF* (neuron-derived neurotrophic factor) in ‘fdqb’ neurons (Fig. 1D), argues for bi-directional and circuit-specific signaling interactions, between motor neurons and muscles, which are likely critical for both their survival (Henderson et al. 1994; Ozaki et al. 2022).

Our muscle samples displayed particularly strong transcriptional segregation according to their anatomical position (Fig. 1C). Although tetrapod limb muscles possess highly distinct morphologies – with stereotypic topology and evolutionarily conserved splitting patterns (Smith-Paredes et al. 2022) – relatively little is known about the molecular differences accompanying this diversity, or their potential relevance for neuromuscular patterning and circuit functionality. Exceptions include the proximal partitioning of early myogenic precursor streams into ventral and dorsal trajectories (Vogel et al. 1996; Schäfer and Braun 1999), or isolated cases of restricted gene activities in a select few limb or limb-adjacent muscles (Haase et al. 2002; Scotti et al. 2015). Here, using genome-wide transcriptional profiling, we substantially expand the list of candidate genes showing muscle-restricted expression profiles (Fig. 1E, Supplementary Table S6). Although limited to three candidate muscles only, these results reveal a molecular complexity in peripheral limb targets that mirrors the diversity observed across distinct LMC motor neuron pools, with likely functional consequences for circuit specificity and refinement (Dasen et al. 2005; Mendelsohn et al. 2017). Of note are the high number of secreted signaling factors and cell surface receptors that drive PCA cluster separation (Fig. 1D,E). Our findings thus extend the framework of molecularly pre-specified identities, established prior or shortly after target contact, from motor neurons to the muscle side of the synapse (Bonanomi and Pfaff 2010; Stifani 2014; Dasen 2017).

To explore this possibility more explicitly, we queried ligand and receptor expression profiles, across our motor neuron and muscle samples, and cross-referenced them with a curated database of cell-cell signaling interactions (Fig. 3) (Ramilowski et al. 2015). Our interaction scoring framework — based on the product of ligand and receptor expression in physically connected neuron-muscle sets — provides a quantitative approach for rapidly screening a large number of candidate pairs, across circuits and in both muscle-nerve and nerve-muscle directions, to prioritize molecular interactions for future functional validation (Fig. 4A,B). The accompanying R Shiny app further enables user-friendly access to these datasets, enabling researchers to explore circuit-specific interaction profiles across all sampled neuron-muscle combinations, in both control and polydactyl configurations, to identify signaling pathways with substantial circuit specificity (Fig. 4C,D). Future studies will need to address the functional significance of the identified candidate pairs, for example through gain- and loss-of-function approaches in the chick embryo or in mouse genetic models, and to determine whether similar molecular address codes operate at other anatomical locations of the neuromuscular system, or in the context of nerve regeneration after injury or disease.

### Transcriptional plasticity and the neuro-muscular axis of limb morphospace exploration

Our analysis of experimentally induced polydactyl limbs further revealed an unexpected degree of transcriptional plasticity. While duplicated muscles largely maintained the molecular identities of their control counterparts, Median nerve-derived motor neurons innervating a duplicated FDQ muscle adopted partial transcriptional features characteristic of canonical FDQ-innervating neurons. This finding suggests that motor neuron identity is not solely determined by intrinsic developmental programs but remains responsive to peripheral target-derived influences. Previous studies have demonstrated that muscle-derived trophic factors such as GDNF regulate motor neuron survival, positioning, and maturation (Haase et al. 2002), while target tissues can influence motor neuron differentiation and connectivity (Bonanomi and Pfaff 2010). Our results extend these observations, suggesting that target muscles may actively reinforce or modify circuit-specific transcriptional programs after initial pathfinding decisions have been made. These findings may also hold relevance for understanding the evolutionary diversification of tetrapod limbs (Dasen 2017).

Over the course of evolution, paired appendages – and particularly the distal autopod, i.e., the hands and feet – have diversified into a vast array of different morphologies, allowing tetrapods to thrive and move on land, in water and in the skies (Rothier et al. 2024). The developmental underpinnings of these changes in structure-function relationships, in particular regarding digit numbers, have so far been predominantly studied at the skeletal level (Zuniga 2015). However, correct integration of skeletal novelties with the neuromuscular apparatus are essential, to ensure limb functionality during morphological transitions (Hirasawa and Kuratani 2018). The ability of muscles to readily attach to the nearest skeletal element offers a highly flexible solution for the limb musculature (Diogo et al. 2015; Luxey et al. 2020, 2023). Additionally, we here show that motor neuron transcriptional identities also remain responsive to peripheral target-derived influences, revealing an unexpected degree of developmental plasticity. Hence, cross-tissue feedback, between muscles and nerves, may facilitate changes in overall limb neuromuscular architecture, thereby enabling motor neurons to adapt to novel muscle topologies in a plastic manner. This may in turn allow for transitional circuit functionality, before adaptive skeletal alterations become evolutionarily canalized (Smith-Paredes et al. 2021; Uller et al. 2024). More broadly, our findings also hold general implications for the role of cell-cell and cell-ECM interactions during cellular transcriptional refinement, highlighting the multi-layered inputs to consider for cell type and cell state manifestations during tissue maturation.

### Conclusion and Outlook

In summary, our study provides evidence that developing neuromuscular circuits of the tetrapod limb are characterized by distinct combinations of transcription factors, ligands, receptors, and effector molecules – in motor neurons and muscles – that may function as molecular address codes during circuit assembly and refinement. Together, these findings deepen our understanding of motor neuron-muscle matchmaking in the tetrapod limb and establish a resource for future investigations into the molecular logic of neuromuscular connectivity, with potential relevance for tetrapod limb evolution as well as regenerative strategies aimed at restoring functional innervation following injury or disease.

## Materials and Methods

### Single motor neuron sampling

Chicken embryos were incubated at 38.5 °C and 59% humidity for 9 days. Polydactyly was induced as previously described and only limbs with 32123 and 321123 digit formulas were considered (Tickle et al. 1982; Pickering, Wali, and Towers 2017; Luxey et al. 2020; Berki et al. 2023). The protocols for single neuron isolation and sequencing have been previously described in (Berki et al. 2023). Briefly, neurons were labelled by retrograde axonal backfill, where a red fluorescent tracer (Alexa Fluor 555 cholera toxin subunit conjugate B, Invitrogen) was injected into the target muscle. After spinal cord dissection and tissue dissociation, the cell suspension was plated in small drops, and fluorescent cells were selected for sequencing after several PBS washes. cDNA quality was assessed on a FragmentAnalyzer instrument (BRAND), and Illumina Nextera XT libraries were prepared on selected high-quality samples (cDNA peak at 1650 to 1700bd, no primer-dimer peak) (Berki et al. 2023). All libraries were sequenced on an Illumina Novaseq 6000 SP sequencer (Median depth: 3,775,397, min/max: 1,974,435/ 21,858,600).

### Muscle bulk sampling

Chicken embryos were incubated at 38.5 °C and 59% humidity for 9 days. Polydactyly was induced as described above. Limbs were separated from the embryo, and individual muscles were dissected and immediately put in RNAlater for RNA extraction. Control muscles were dissected from the non-duplicated limb of the treated embryos. Libraries were prepared for sequencing according to the SMARTSEQ2 GOTTGEN protocol and paired-end sequenced for 51 cycles on an Illumina NovaSeq 6000 sequencer (Median depth: 29,176,887, min/max: 23,555,023/ 37,294,941).

### Sequence data analysis

The same custom pipeline was used for muscle and neuron data to map the raw reads to the GRCg6a chicken genome. Briefly, the steps include: (1) trimming reads for low-quality sequences and sequencing adapters with Trimmomatic v.0.39 (Bolger, Lohse, and Usadel 2014); (2) read alignment against *Galgal6* chicken genome (https://ncbi.nlm.nih.gov/datasets/genome/GCA_000002315.4/) with STAR aligner v.2.5.2 (Dobin et al. 2013); (3) creating count tables with HTSeq v.0.6.1 (Anders, Pyl, and Huber 2015). Transcript Per Million (TPM) tables were created with StringTie v.2.1.0 (Pertea et al. 2015). All subsequent analyses were conducted in R-4.4.2 (R Core Team 2024).

### Muscle data analysis

Raw count tables were transformed into a DESeqDataSet (dds) with the R package DESeq2, filtering out W chromosome genes (Love, Huber, and Anders 2014). Muscles were filtered by number of genes detected (nGenes > 14,000), expression of myosin (*MYL1, MYL2*, and *MYL10*), troponin (*TNNI1, TNNI2, TNNC1*, and *TNNC2*) and other muscle markers (*MYOD1* and *MYOG*) and inspection by principal component analysis (PCA) to identify outliers. Differential expression (DE) analyses were performed in DESeq2, using the function DESeq() and results() (adjusted *p-value* < 0.05), on dds subsets for the compared muscles.

### Neuron data analysis

In addition to DESeq2, we used zinbwave, an R package tailored to scRNA-seq data, to create a DESeqDataSet (Love et al. 2014; Risso et al. 2018). We followed the packages vignettes for data transformation and normalization. First, observational weights were calculated with zinbwave() (K = 0, epsilon = 1e12) and then transformed using DESeqDataSet(). Then, size factors were calculated with scran’s computeSumFactors() before running DESeq() (test = “LRT”, reduced = ∼1, minmu = 1e-6, minReplicatesForReplace = Inf). In addition, motor neurons were filtered for percentage of mitochondrial RNA (perc_mt < 0.25), hemoglobin expression (*HBA1, HBAD, HBBR, HBZ, HBE, HBE1*), motor neuron marker expression (*ALDH1A2, FOXP1, SLC18A3*, and *CHAT*) and inspection by PCA to identify low-quality outliers. Visualizations of neuron data analyses were performed similarly to the muscle data.

### Data visualization

PCAs were created with the function mPCA from the R package modplots (https://github.com/safabio/modplots), which uses DESeq2’s plotPCA function, using the top 500 most variable genes (Love et al. 2014; safabio [2022] 2025). Z-scored heatmaps of DE genes were obtained with mPheatmapDESeq2 from the modplots R package. Hheatmaps based on log1p-transformed expression and Pearson correlations were generated with pheatmap() (Kolde 2019). Volcano plots were obtained using the R package ggplot2 (Ginestet 2011). Gene Ontology (GO) Term enrichment analyses were conducted on DE genes using the function goana from the limma R package (Ritchie et al. 2015), and visualized using ggplot2 (Ginestet 2011). Finally, upset plots were created with the UpSetR package to visualize intersections of various sets of DE genes (Gehlenborg 2019).

### Immunohistochemistry

Polydactyl embryos were dissected at day 10. Polydactyl limbs were fixed in 4% PFA overnight, cryoprotected in 30% Sucrose solution, then embedded in blocks (FSC22 Clear Frozen Section Compound, Biosystems) and kept at -80°C until sectioning. 20 μm thick sections were cut and thawed for 15 min, washed in PBS to eliminate OCT and blocked in PBST (0.1% Triton, 0.02% SDS, 0.002% BSA in PBS) solution for an hour. Primary mouse antibodies MF20 (DSHB) and Na8 (DSHB) were diluted 1/500 in PBST and adjacent slides were incubated with the mix in a humid chamber overnight at 4 °C. After three PBST washes, anti-mouse Cy3 secondary antibodies (1/500) were deposited on the slides, incubated for 2 hours at room-temperature, washed three times in PBST and mounted with fluorescent mounting media (Agilent). All slides were treated with DAPI for 15 minutes in PBS, then washed and mounted. Slides were kept at 4 °C until imaging with an Olympus Fluoview FV3000 confocal microscope using a 10x/0.4 objective (air, ApoPlan, Olympus).

### Interaction scores

To calculate interaction scores, the ligand-receptor interaction table from Ramilowski et al. 2015 was filtered for interactions with chicken orthologs expressed in our data set (Ramilowski et al. 2015). The interactions were scored as the product of TPM values for the ligand and the receptor. To screen for relevant interactions, scores were calculated for ‘correct’ (corresponding neuron muscle pairs) and ‘incorrect’ (non-corresponding neuron muscle pairs) interactions, calculating p-values with rstatix’s wilcox_test() (alternative = “greater”, p.adjust.method = “BH”) to highlight specific interactions (Kassambara 2023). Correct interactions were defined as following: FCU-fcu_(dup)_, AM-am_(dup)_, FDQ-fdq_a_, FDQ-fdq_b_, FDQ_dup_-fdq_dup_ (Muscle ligand-neuron receptor); fcu_(dup)_-FCU, am_(dup)_-AM, fdq_a_-FDQ, fdq_b_-FDQ, fdq_dup_-FDQ_dup_ (Neuron ligand-muscle receptor). Plots were created with ggplot2 (Ginestet 2011). Due to a limited number of FCU and AM connecting neurons, we pooled control and duplicated muscle innervating samples and disregarded interactions with FCU_dup_ and AM_dup_ muscles altogether.

### Interactive app

The interactive app was built using R Shiny. The app was then dockerized using the shiny2docker package. Docker build and push to ghcr.io is handled by a github actions workflow https://github.com/safabio/crosstalk_paper_shiny

## Code and App availability

Code used for analysis and figure generation is accessible under https://github.com/safabio/crosstalk_paper The Shiny app is available as a docker container hosted on https://ghcr.io/safabio/crosstalk_paper_shiny

## Acknowledgments

The authors wish to thank E. Stoeckli and all members of the group for useful discussions. Calculations for scRNA-seq analyses were performed at sciCORE (http://scicore.unibas.ch/), scientific computing center at the University of Basel. This work was supported by funds from the Janggen-Pöhn Foundation to F.S., and the Swiss National Science Foundation (SNSF project grant 310030_170022), the Olga Mayenfisch Foundation, the FSRMM, and the University of Basel to P.T.

## Supplementary Figures

**Figure S1. (related to Figure 1).**
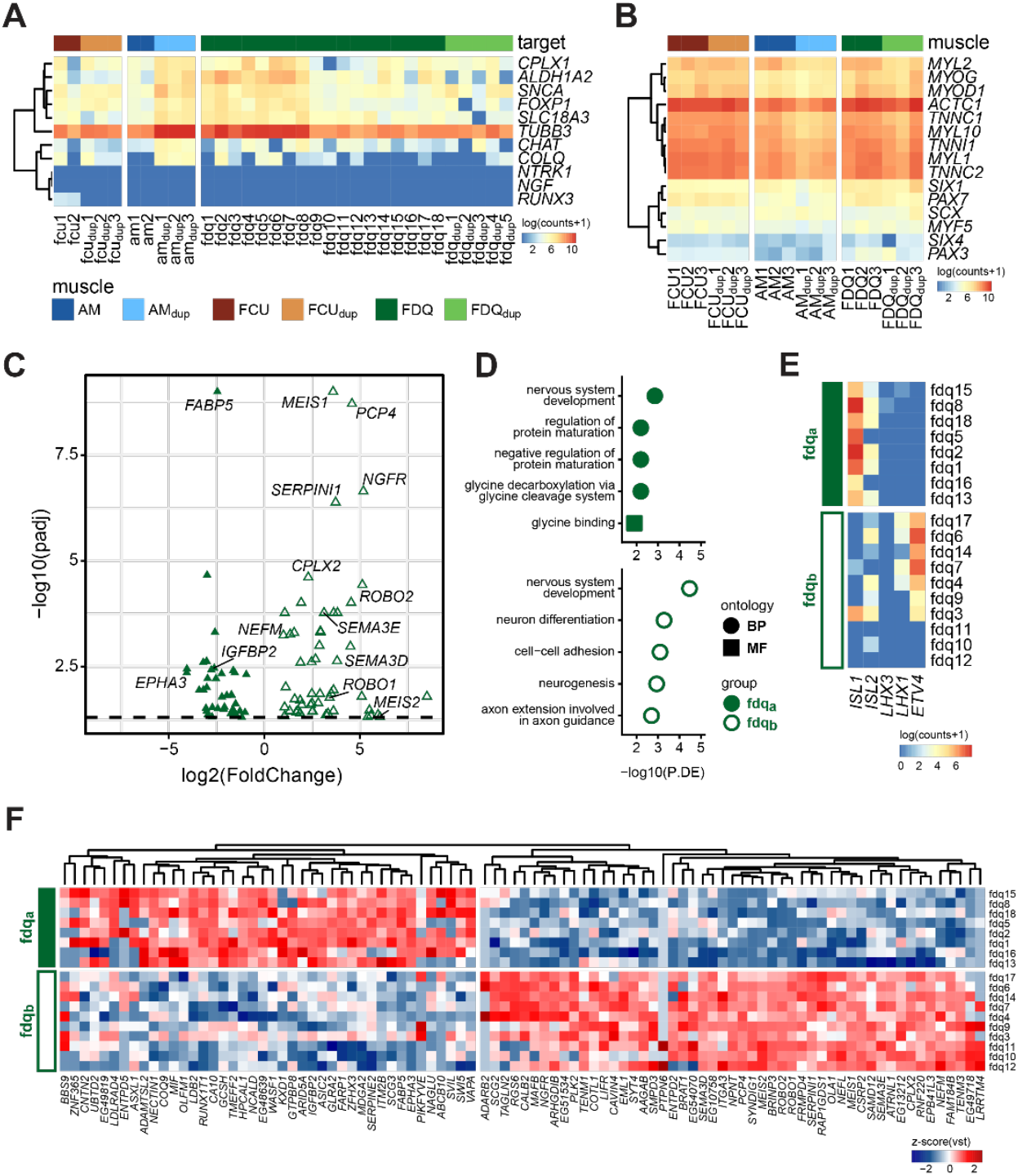
**(A)** Expression heatmap of canonical markers for neurons (*TUBB3*), sensory neurons (*NTRK1, NGF, and RUNX3)*, and motor neurons (*CPLX1, ALDH1A2, SNCA, FOXP1, SLC18A3, CHAT, and COLQ*). Samples are colored by their target muscle. **(B)** Heatmap of marker gene expression for muscle samples. Colors correspond to Fig. 2A. **(C)** Volcano plot showing differential expression of DE genes between neurons belonging to ‘fdq_a_’ and ‘fdq_b_’. Maximum -log_10_(p-adj) value displayed is capped at 9 (*FABP5* and *MEIS1*) **(D)** Selected gene ontology terms enriched in the DE genes between neurons belonging to FDQ_a_ and FDQ_b_. **(E)** Expression heatmap of motor column markers, for MMC (*LHX3, ISL1, ISL2*), LMC_m_ (*ISL1, ISL2*), and LMC_l_ (*LHX1, ISL2, ETV4*). **(F)** Heatmap of z-scored variance stabilizing transformation expression values for differentially expressed genes between neurons belonging to ‘fdq_a_’ and ‘fdq_b_’, with an adjusted p-value < 0.05.

**Figure S2. (related to Figure 2).**
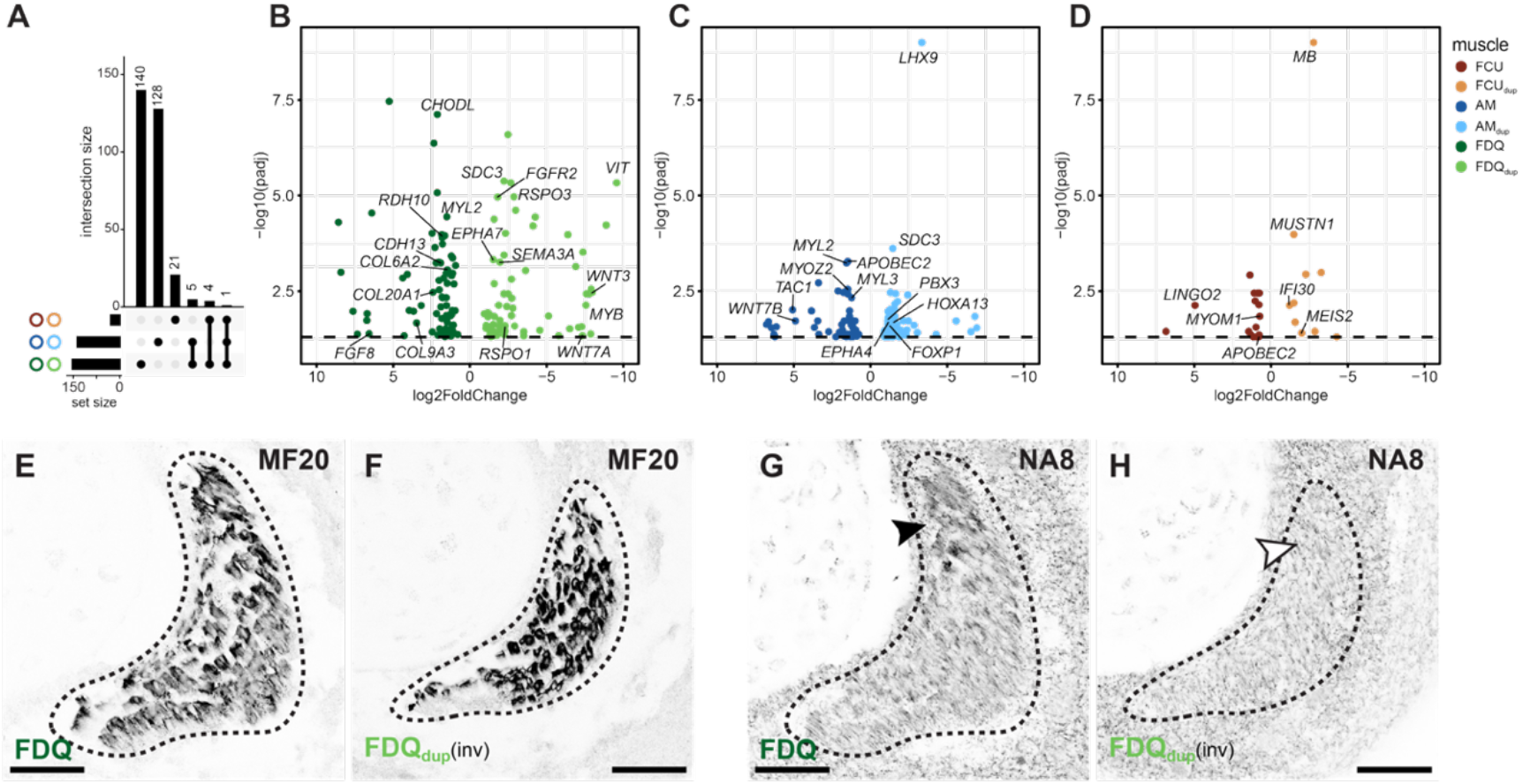
**(A)** Upset plot showing the intersections of differentially expressed (DE) genes between control and duplicated muscles. **(B-D)** Volcano plots of DE genes between control muscles and their duplicated counterparts. Maximum -log10(p-adj) value displayed is capped at 9 (*LHX9* and *MB*). **(E-H)** Immunohistochemistry for MF20 (muscle specific myosin heavy chain) and NA8 (slow twitch muscle fibres) on FDQ and FDQdup muscles from a wing with induced polydactyly. Brightness and contrast were adjusted globally, and images of duplicated FDQdup muscles were inverted, for better comparison. Scale bars are 50 µm.

**Figure S3. (related to Figure 2).**
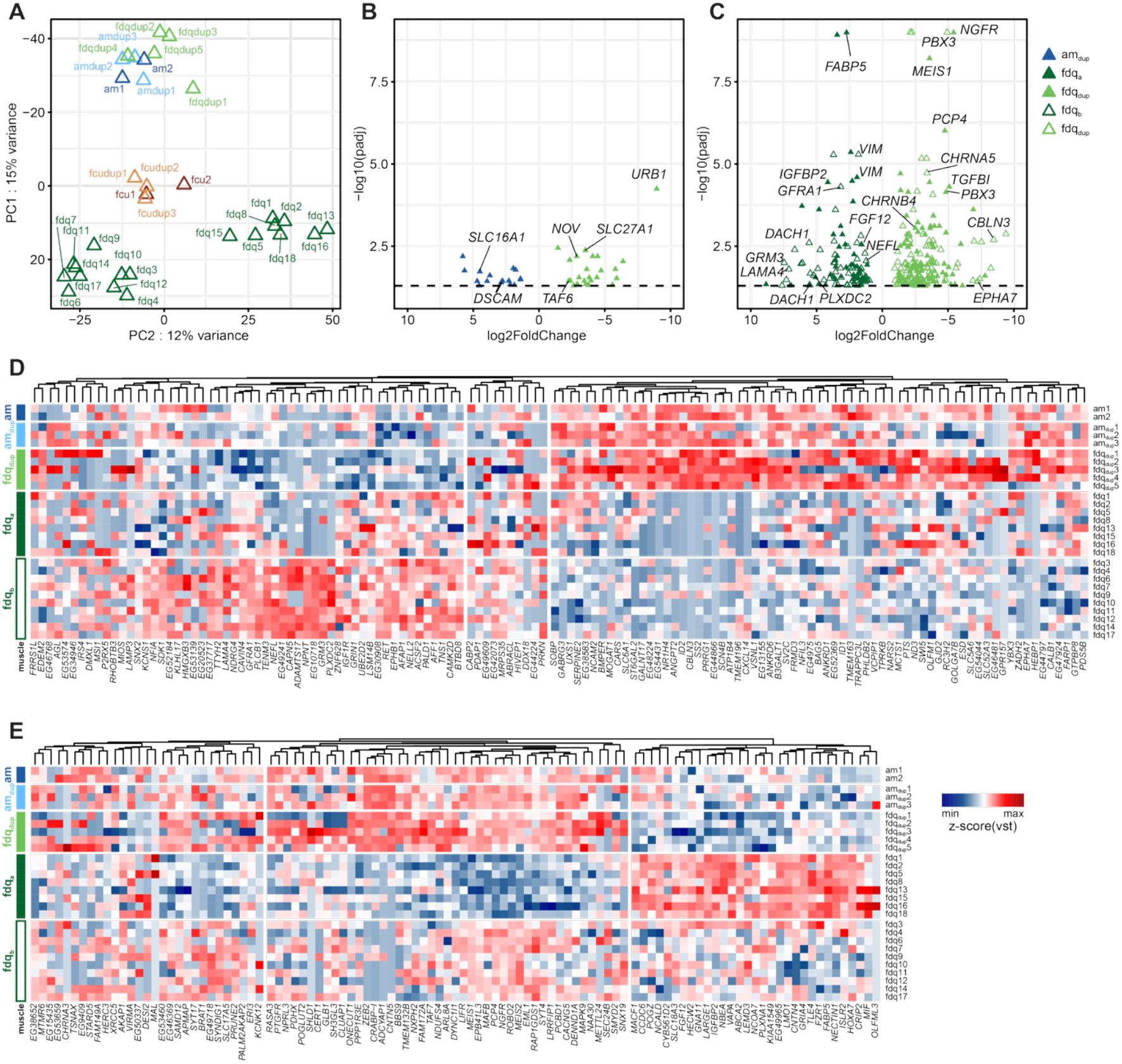
**(A)** PC1 and 2 of the PCA of all neurons based on the top 500 variable genes. Axes are arranged to correspond to Fig. 2B. Colors correspond to Fig. 2A. **(B)** Volcano plot showing differentially expressed genes between neurons belonging to ‘am_dup_’ and ‘fdq_dup_’. **(C)** Volcano plot showing differentially expressed genes, from ‘fdq_a_’ versus ‘fdq_dup_’ contrasts (full triangles) and ‘fdq_b_’ versus ‘fdq_dup_’ contrasts (empty triangles). Maximum -log10(p-adj) value displayed is capped at 9. **(D-E)** Heatmap of z-scored vst expression values for the DE genes between ‘fdq_b_’ and ‘fdq_dup_’ **(D)** or ‘fdq_a_’ and ‘fdq_dup_’ neurons **(E)**. Columns clustered by Euclidean distance. Rows are ordered by target muscle.

**Figure S4. (related to Figure 3).**
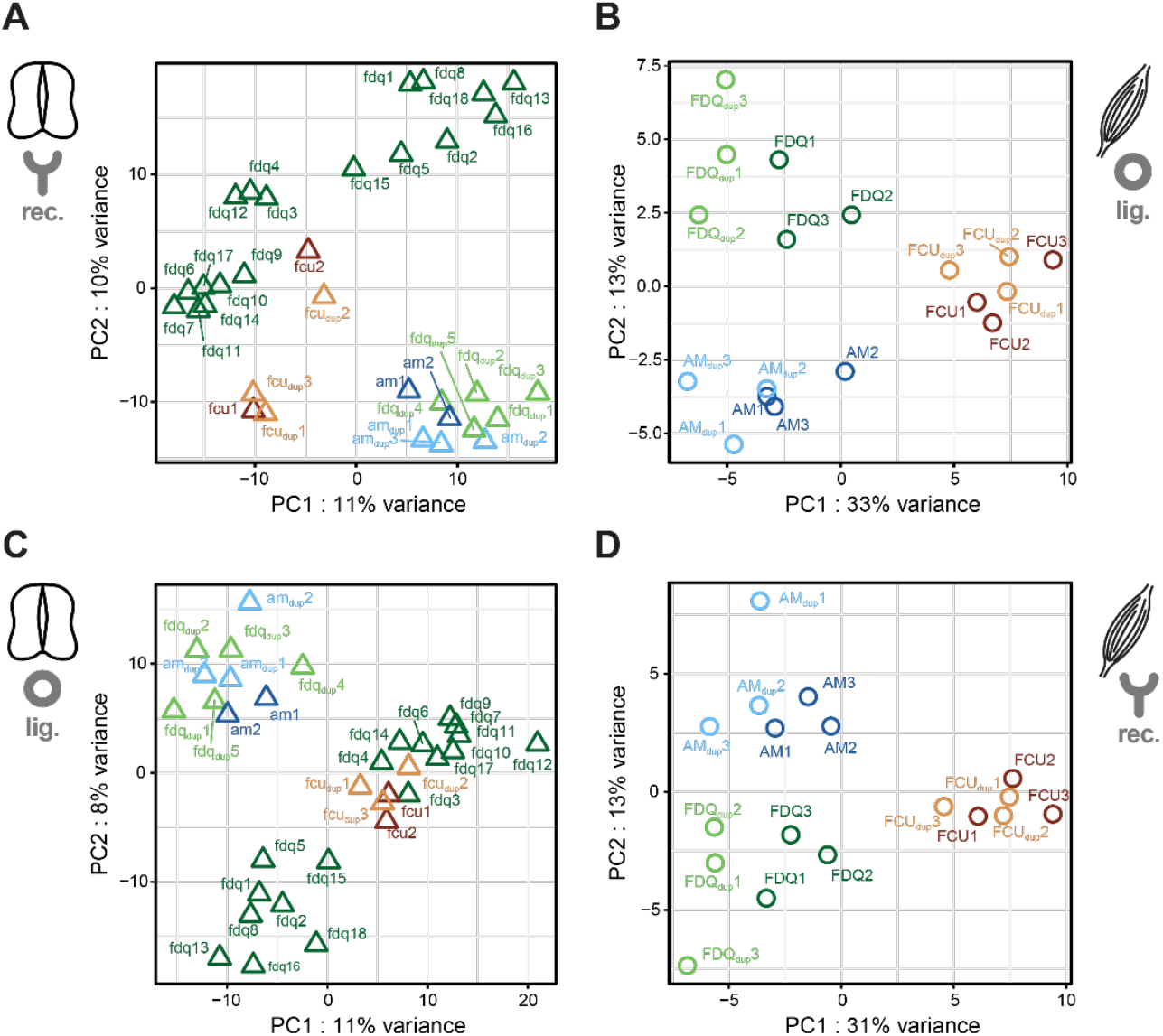
**(A-D)** PCA based on expression profiles of ligand and receptors. **(A)** Receptors expressed in neurons (n = 379). **(B)** Ligands expressed in muscles (n = 380). **(C)** Ligands expressed in neurons (n = 307). **(D)** Receptors expressed in muscles (n = 453). PCA = principal component analysis.

